## Supplemental Material for "A polarity pathway for exocyst-dependent intracellular tube extension"

**Figure S1. *t28h11.8p* is an excretory cell-specific promoter during embryonic and larval canal outgrowth**

(A) Widefield fluorescence images of *t28h11.8p::mCherry* (‘*excP::mCh*’) transcriptional reporter during embryonic elongation. 3-fold stage of embryo elongation is shown as this represents the initial stage of posterior canal growth. *t28h11.8p::mCherry* expression could not be visually detected in any tissues outside of the excretory canal during embryogenesis. (B) *t28h11.8p::mCh* expression during the L1 larval stage, as canal growth proceeds beyond half of the animal’s body length. Excretory cell body indicated by asterisk. Posterior tip of excretory canal indicated by white arrow. Outline of each animal is indicated by solid white line. A single canal arm is shown in each image with anterior canal extensions visible adjacent to the cell body. Scale bars, 10 μm. ‘Unsharp mask’ filter was applied equally to all images using ImageJ software.

**Figure S2. The SEC-5$^{exc(-)}$ canal outgrowth phenotype is not enhanced by a *sec-5* null allele**

Canal outgrowth defects upon depleting the indicated proteins in the excretory cell are indicated as the percentage of animals in each of four phenotypic categories that measure posterior canal extension relative to body length (see Figure 2). The relative intensity of green shading reflects the percentage of larvae observed in each phenotypic category. The *P*-value was calculated using Fisher’s exact test (<50% versus >50% canal outgrowth).

**Figure S3. CDC-42 depletion causes a split lumen phenotype in larval excretory canals**

(A-B'') Widefield fluorescence images of larval excretory canal phenotypes in CDC-$42^{exc(-)}$ L1 and L4 larval stage worms expressing cytoplasmic and lumenal markers. An additional lumen that has split off of the canal arm is indicated by white arrowhead. Scale bars, 10 μm. 'Unsharp mask' filter was applied equally to all images using ImageJ software.

**Figure S4. Depletion of PAR-3 causes mild excretory cell lumen defects during early larval stages**

(A-B'') Larval excretory canal phenotypes in PAR-$3^{exc(-)}$ L1 stage worms expressing cytoplasmic and lumenal markers. Images are of the same animal at different magnifications, 20x (A) and 63x (B-B''). Excretory cell body indicated by asterisk. Posterior tip of excretory canal indicated by white arrow. Outline of animal is indicated by solid white line. Single canal arm is shown in each image with anterior canal extensions visible adjacent to cell body. Scale bars, 10 μm.

**Figure S5. PAR-6::ZF1::YFP depletion by acute ZIF-1 expression**

(A-B) Distribution of PAR-6::ZF1::YFP in larval excretory canal in control (A) and *hspP::zif-1* (B). (C) Quantification of PAR-6::ZF1::YFP intensity in the excretory canal of control and *hspP::zif-1* larvae. Individual data points from a single experiment are represented by black dots, horizontal bar is the mean, and error bars are the SEM. Outline of excretory canal cytoplasm is indicated by dotted line. Scale bar, 10 μm.

Figure S1

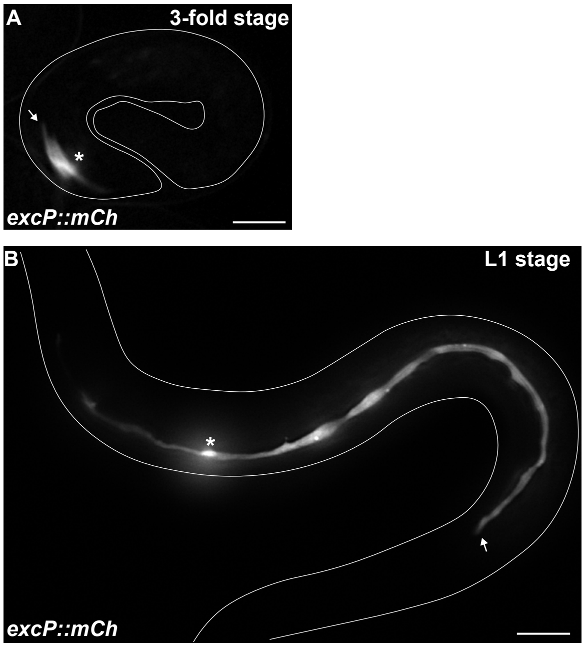

### Figure S2

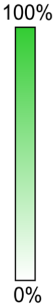


L4 stage

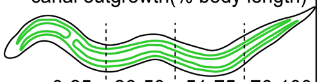

| | canal outgrowth(% body length) | | | | | |
|---|---|---|---|---|---|---|
| | 0-25 | 26-50 | 51-75 | 76-100 | *n* | *P* value |
| SEC-5$^{exc(-)}$ | 62 | 38 | 0 | 0 | 21 | |
| SEC-5$^{exc(-)}$/*sec-5(tm1443)* | 61 | 29 | 10 | 0 | 49 | 0.3127 |

### Figure S3

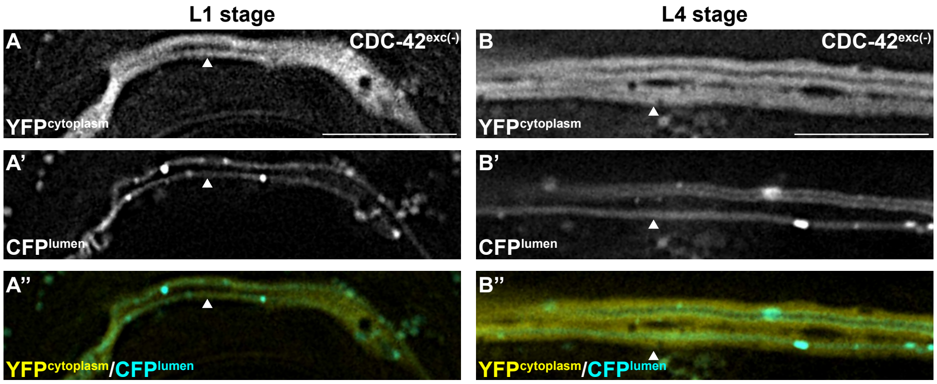
L1 stage
L4 stage
A
CDC-42exc(-)
YFPcytoplasm
B
CDC-42exc(-)
YFPcytoplasm
A'
CFPlumen
B'
CFPlumen
A''
YFPcytoplasm/CFPlumen
B''
YFPcytoplasm/CFPlumen

Figure S4

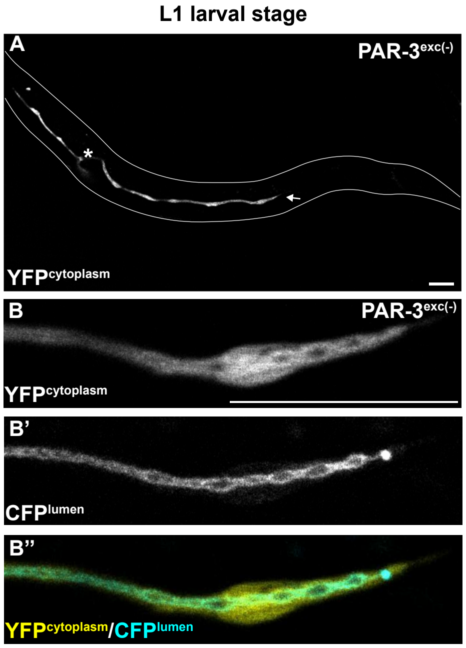
L1 larval stage
A
PAR-3exc(-)
*
YFPcytoplasm
B
PAR-3exc(-)
YFPcytoplasm
B'
CFPlumen
B"
YFPcytoplasm/CFPlumen

Figure S5

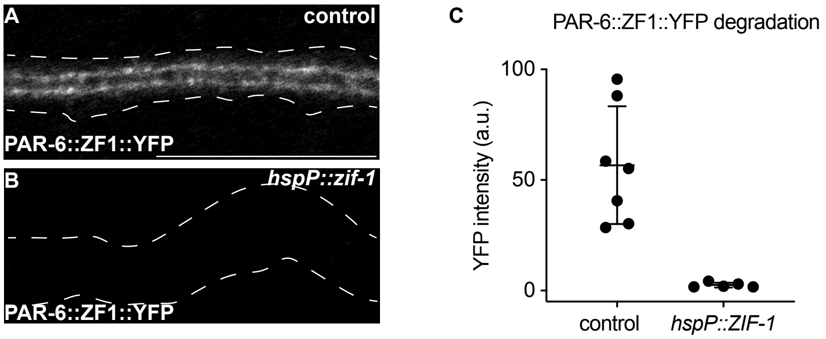
A
control
PAR-6::ZF1::YFP
B
hspP::zif-1
PAR-6::ZF1::YFP
C
PAR-6::ZF1::YFP degradation
100
50
0
YFP intensity (a.u.)
control
hspP::ZIF-1

Table S1. ***C. elegans strains***

| Strain | Genotype |
|---|---|
| N2 | wild type |
| FT95 | *xnIs23[cdc-42p::zf1::gfp::cdc-42 unc-119(+)]; unc-119(ed3)* |
| FT1202 | *sec-5(tm1443)/mIn1[mIs14 dpy-10(e128)]* |
| FT1452 | *xnSi37[pgp-12p::mCherry unc-119(+)]; unc-119(ed3)* |
| FT1523 | *sec-5(xn51[sec-5::zf1::yfp loxP unc-119(+) loxP]); unc-119(ed3)* |
| FT1692 | *xnIs23; xnEx437[t28h11.8p::mCherry, t28h11.8p::zif-1]; unc-119(ed3)* |
| FT1699 | *par-3(xn59[par-3::zf1::yfp loxP unc-119(+) loxP]); unc-119(ed3)* |
| FT1702 | *par-6(xn60[par-6::zf1::yfp loxP unc-119(+) loxP]); unc-119(ed3)* |
| FT1834 | *sec-5(xn51); xnIs547[t28h11.8p::zif-1]; par-3(it301[par-3::mCherry]); xnEx466[t28h11.8p::yfp::sl2::ifb-1::cfp, pRF4]* |
| FT1837 | *xnIs547; par-3(it301); xnEx466* |
| FT1844 | *par-6(xn60); xnIs547; xnSi31[sec-8p::sec-8::mCherry unc-119(+)]; xnEx473[t28h11.8p::yfp::sl2::ifb-1::cfp, pRF4]* |
| FT1846 | *par-3(xn59); xnIs547; xnSi31; xnEx475[t28h11.8p::yfp::sl2::ifb-1::cfp, pRF4]* |
| FT1849 | *cdc-42(xn65[zf1::yfp::cdc-42 loxP unc-119(+) loxP]); xnIs547; par-3(it301); xnEx477[t28h11.8p::yfp::sl2::ifb-1::cfp, pRF4]* |
| FT1866 | *ral-1(tm5205); xnIs472[ral-1p::zf1::yfp::ral-1]; xnIs547; xnEx472[t28h11.8p::yfp::sl2::ifb-1::cfp, pRF4]* |
| FT1942 | *pkc-3(xn84[zf1::gfp::pkc-3]); xnIs547; xnEx466* |
| FT1945 | *cdc-42(xn65); par-6(cp60[par-6::mKate::3xMyc loxP unc-119(+) loxP]); xnEx481[hsp-16.41p::zif-1; t28h11.8p::yfp::sl2::ifb-1::cfp, pRF4]* |
| FT2015 | *par-3(xn59); par-6(cp60); xnEx491[t28h11.8p::cfp, pRF4]* |
| FT2020 | *par-6(xn60); par-3(it301); xnEx494[hsp-16.41p::zif-1; t28h11.8p::CFP, pRF4]* |
| FT2022 | *par-6(xn60); par-3(it301); xnEx496[t28h11.8p::CFP, pRF4]* |
| FT2027 | *par-3(xn59); par-6(cp60); xnEx501[hsp-16.41p::zif-1; t28h11.8p::CFP, pRF4]* |
| FT2061 | *par-6(xn60); xnIs485[sec-10p::mCherry::sec-10]; xnEx508[hsp-16.41p::zif-1; t28h11.8p::CFP, pRF4]* |
| FT2065 | *par-6(xn60); xnIs485; xnEx511[t28h11.8p::cfp, pRF4]* |
| FT2069 | *par-3(xn59); xnIs485; xnEx514[t28h11.8p::cfp, pRF4]* |
| FT2089 | *exc-5(xn108[exc-5::zf1::mScarlet]); pkc-3(it309[gfp::pkc-3]); xnEx519[hsp-16.41p::zif-1; t28h11.8p::CFP, pRF4]* |
| FT2093 | *exc-5(xn108); pkc-3(it309); xnEx523[t28h11.8p::cfp, pRF4]* |
| FT2100 | *par-3(xn59); xnIs485; xnEx528[hsp-16.41p::zif-1; t28h11.8p::CFP, pRF4]* |
| FT2289 | *cdc-42(xn65); par-6(cp60); xnEx551[hsp-16.41p::zif-1; t28h11.8p::CFP, pRF4]* |
| KK1218 | *par-3(it301)* |
| KK1228 | *pkc-3(it309)* |
| LP282 | *par-6(cp60); par-3(cp54[mNeonGreen::3xFlag::par-3])* |
